# Distinct forms of amyloid-β moderate sleep duration through NAD^+^-linked redox metabolism in Alzheimer’s disease

**DOI:** 10.1101/2025.02.26.640405

**Authors:** Yizhou Yu, Giorgio Fedele, Ivana Celardo, Lilian Zhou, Bryan Wei Zhi Tan, Samantha H. Y. Loh, L. Miguel Martins

## Abstract

Sleep disruptions precede a clinical diagnosis of Alzheimer’s disease (AD) by several years. However, how AD pathologies affect sleep remains unclear. Here, we integrate epidemiological data with insights from *Drosophila* models of AD to investigate how AD progression could be linked to sleep disruption. We found that individuals with a high risk of AD report a shorter sleep duration than do those with a clinical diagnosis of AD. We showed that the expression of different forms of amyloid-β in flies can replicate these variations in sleep duration. Analysis of the metabolome and proteome of these flies revealed distinct changes in NAD^+^-linked redox metabolism and levels of hyperkinetic (Hk), a redox-sensing sleep homeostat. We showed that the genetic upregulation of Hk is neuroprotective and increased *KCNAB2* expression in the brain decreases AD risk in humans. Overall, our data provides a new mechanism linking the disruption of NAD^+^-linked redox sensing and sleep disruption in AD.

## Introduction

Alzheimer’s disease (AD) is the most prevalent neurodegenerative disease worldwide and is projected to affect more than 150 million people globally by 2050 (Nichols et al., 2022). Molecular hallmarks of AD, such as the accumulation of amyloid-β (Aβ) and tau aggregates, are thought to precede neurodegeneration (Alzheimer, 1907; Bird, 1993). Increases in the levels of Aβ occur decades before disease onset and cause neuronal toxicity. These molecular hallmarks, combined with genetic risk, age, and social and physiological health determinants, contribute to AD risk (Livingston et al., 2020).

Sleep disruption is a risk factor for AD, and poor sleep often precedes a clinical diagnosis (Ju et al., 2014). However, studies on the causal relationship between sleep and AD remain contradictory. Some studies suggest that shorter sleep duration (insomnia) increases AD risk (Sabia et al., 2021). A lack of sleep is linked to increased neuronal excitability (Tabuchi et al., 2015) and increased Aβ levels (Shokri-Kojori et al., 2018). However, other studies suggest that longer sleep durations (hypersomnia) can exacerbate AD pathologies. A recent study proposed that reducing sleep duration could be used as a therapy for AD (P. Li et al., 2022), but reducing sleep duration can disrupt neuronal function (Acosta-peña et al., 2015). Sleep duration and brain health have a nonlinear relationship, with deviations from 7 hours of sleep linked to poorer cognitive function (Y. Li et al., 2022).

The fruit fly *Drosophila melanogaster* is a powerful model for studying the mechanisms of neurodegeneration (Muqit and Feany, 2002) and sleep (Sehgal and Mignot, 2011). Sleep integrates circadian (Konopka and Benzer, 1971; Vosshall et al., 1994; Zehring et al., 1984) and homeostatic (Borbély, 1982) components, which are conserved from flies to mammals (Allada et al., 2017). The homeostatic component of fly sleep has similar molecular and behavioural correlation to that in mammals (Hendricks et al., 2001; Shaw et al., 2000). The homeostatic sleep switch in flies integrates metabolic information to modulate neuronal function (Kempf et al., 2019; Pimentel et al., 2016). Hyperkinetic (Hk), a potassium channel subunit, senses NADP^+^ and NADPH levels to alter the firing properties of sleep neurons (Kempf et al., 2019). Loss of Hk impairs sleep and cognition (Bushey et al., 2007), but the connection between Hk and AD is yet unknown.

*Drosophila* models of AD are created by expressing human proteins in the fly brain (Crowther et al., 2005; Luheshi et al., 2007; Moloney et al., 2010). These models exhibit conserved features of AD pathologies in human patients, ranging from cellular dysfunction, such as mitochondrial impairment (Yu et al., 2021a), to behavioural deficits, such as memory loss (Wang et al., 2012).

AD is associated with increased Aβ aggregation in the brain (Selkoe, 1991; Selkoe and Hardy, 2016). Administering antibodies to AD patients to eliminate Aβ delays cognitive decline (Sims et al., 2023; van Dyck et al., 2023), indicating that Aβ is a causal factor in AD pathology. Individual monomers of Aβ self-assemble to form oligomers, protofibrils and larger fibrils (reviewed in (Hampel et al., 2021; Scheres et al., 2023). The degree of aggregation of Aβ is positively correlated with its ability to decrease lifespan in AD models. The Arctic mutation (Aβ42Arc), a specific variant of Aβ42, exemplifies this as it tends to form protofibrils, a critical step in the pathogenic process (Nilsberth et al., 2001; Yang et al., 2023). This mutation, characterised by a glutamate to glycine substitution at position 22, enhances the nucleation phase of fibril formation, thus accelerating the aggregation process in neuronal tissue. The increased aggregation of Aβ42Arc is linked to more rapid cognitive decline in AD patients and increased neurotoxicity in fly models of AD (Moloney et al., 2010; Yu et al., 2021a).

Here, we combine medical data from AD patients with experimental work using a fly model of AD involving the expression of two different forms of Aβ to uncover potential mechanisms of sleep disruption in AD. Using fly models and medical records from UK Biobank participants, we found that individuals at risk of AD have a decreased sleep duration, whereas those with an AD diagnosis sleep more. In fly models of AD, the expression of monomeric Aβ caused a decrease in sleep duration, mimicking the decreased sleep duration in at-risk individuals. Expressing an aggregate-prone form of Aβ increased sleep duration, which is consistent with the increased sleep duration in AD patients with a clinical diagnosis. To explore the complex relationship between sleep and AD at the molecular level, we analysed global changes in the metabolome and proteome of flies that have a divergent sleep phenotype caused by the expression of the monomeric and aggregate-prone forms of Aβ. We found that the protein levels of Hk were different in these fly models of AD and that the overexpression of Hk increased the lifespan of flies expressing aggregate-prone Aβ. We found that neurons from post-mortem donors with severe AD also have increased expression of *KCNAB*, the orthologue of *Hk*. Through Mendelian randomisation, we found that increased *KCNAB2* expression in neurons is associated with decreased AD risk in humans. Taken together, our work provides mechanistic insights into the complex relationship between sleep and AD through *Hk*/*KCNNAB2*.

## Results

### Alzheimer’s disease progression is linked to distinct changes in sleep duration

Age and age-related diseases, including AD, are linked to complex changes in sleep duration. Sleep duration forms a U-shaped trend across age in humans, with the shortest sleep duration observed in adults aged between 35 and 55 years (Coutrot et al., 2022). Sleep disturbances, such as shortened or fragmented sleep, are both early symptoms of AD and potential contributors to its progression (Minakawa et al., 2019). However, the molecular mechanisms behind the age-related changes in sleep duration and how these changes might be connected with AD remain unclear. AD pathology progresses through a series of pathological changes and clinical manifestations (McKhann et al., 2011). At an early preclinical stage, AD has no symptoms, followed by a middle stage in which AD patients have mild cognitive impairment. Finally, patients reach a clinical stage marked by clinical symptoms of dementia. To explore the links between sleep duration and different AD stages, we analysed medical data from the UK Biobank (Figure 1a). We observed a general trend in which younger AD patients had a shorter sleep duration than did individuals without AD. Older AD patients reported longer sleep durations than did individuals without an AD diagnosis (Figure 1b). We hypothesised that sleep duration is linked to AD stage. First, we modelled a preclinical stage of AD using a polygenic risk score (PRS) of AD. The heritability of AD was estimated at 58% (Gatz et al., 2006), and this PRS is linked to an increased risk of AD (Yu et al., 2021a). We used this PRS to mimic the preclinical stage of AD (schematic in Figure 1c). We found that a high genetic risk of AD was not associated with changes in daytime sleepiness (Figure 1d) but was associated with a significant reduction in nighttime sleep duration (Figure 1e) after adjusting for covariates such as age and sex. Next, we investigated the sleep duration of participants with a clinical diagnosis of AD. We found that a clinical diagnosis of AD was linked to both a significantly greater level of daytime sleepiness (Figure 1f) and nighttime sleep duration (Figure 1g). We thus conclude that AD progression exacerbates age-related changes in sleep duration.

**Figure 1.**
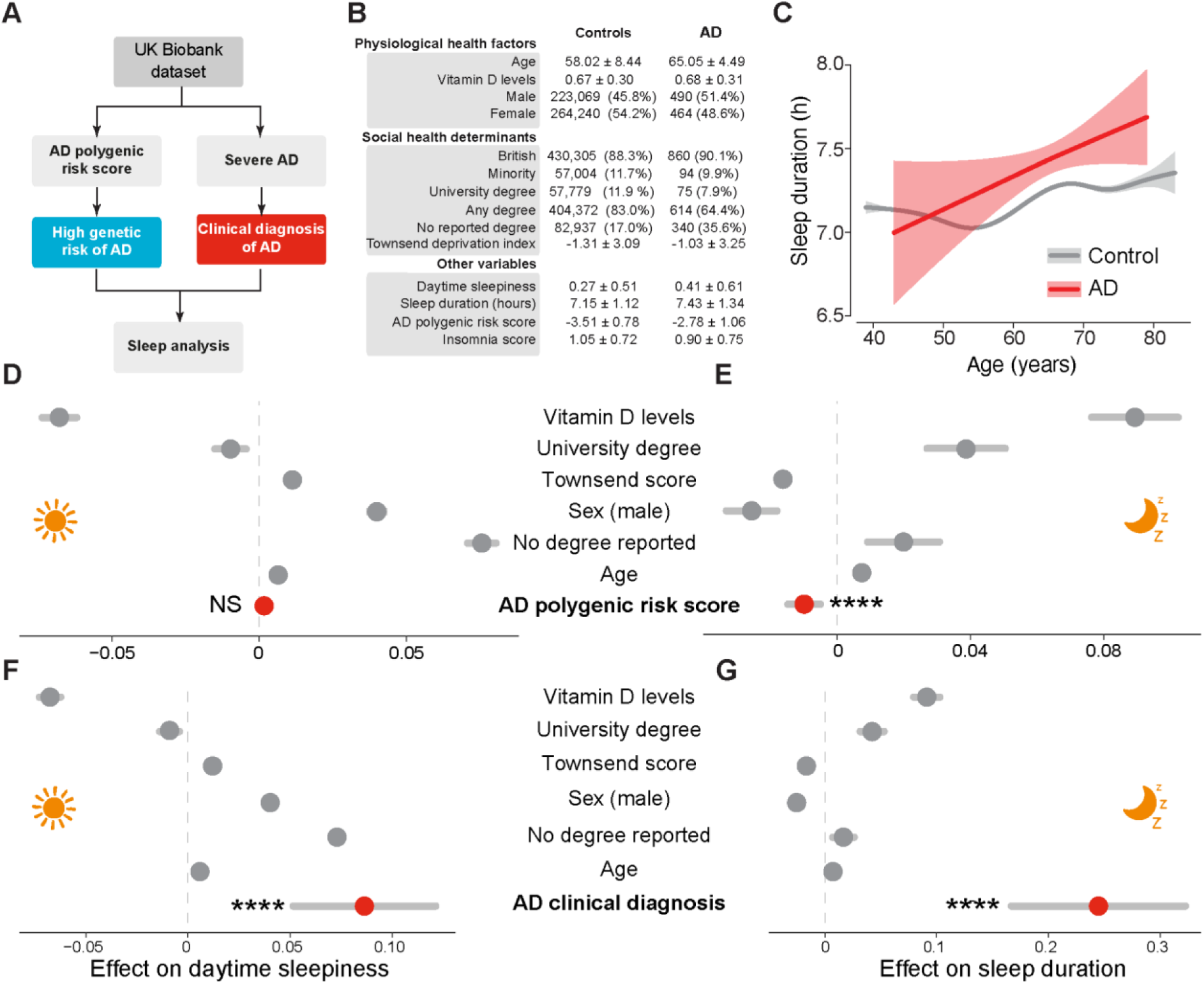
Participants with high AD risk reported short sleep durations, whereas AD patients reported longer sleep durations. **A**. Descriptive statistics of the UK Biobank cohort analysed. Either the number of participants in each category and their percentage with respect to the total cohort or the mean and standard deviation are shown. **B**. Changes in self-reported sleep duration across age in patients with an AD diagnosis compared with healthy controls across age. Generalised additive models were used to fit the data for both control and AD patients. The trendline is in red for AD patients and grey for controls. The standard errors are in the same colours but with decreased opacity. **C**. Rational in separating individuals with high and low AD pathology. A clinical diagnosis of AD (right arm) is based on cognitive impairments, which occur at a later stage of AD. We thus refer to individuals with a diagnosis of AD as those with “severe” AD. To model a preclinical model of AD, we applied a polygenic risk score in individuals without a diagnosis of AD. Individuals with a higher genetic risk of AD would have an increased AD burden. **D** and **E**. UK Biobank participants with a high genetic risk of developing AD do not experience daytime dozing (**D**) but report sleeping less (**E**) during the night (linear regression, asterisk). **F** and **G**. Patients with a clinical diagnosis of AD experience increased daytime sleepiness (**F**, linear regression, asterisks) and longer sleep duration at night (**G**, linear regression, asterisks). For **C–F**, confounding variables are labelled in grey, the targeted variable is labelled in red, and their respective 95% confidence intervals are labelled in grey. Statistical significance through asterisks is only indicated for the investigated variable for clarity.

These observations might be linked to multiple factors, including biological, societal and psychological. To investigate the causal effects of AD progression on sleep duration in a controlled environment, we used a fruit fly model of AD that expresses Aβ42Arc. First, we found an age-related increase in sleep duration in flies, similar to AD patients with a clinical diagnosis (Figure 2a). Compared with matched controls, young flies expressing Aβ42Arc have shorter sleep durations during the dark phase (Figure 2b), similar to the preclinical model of AD. We next analysed how the sleep duration changes during the light and dark phases across the lifespan of Aβ-expressing flies compared with controls. We observed a similar decrease in sleep duration at a younger age, with an increase in sleep duration related to age (Figure 2c), mimicking our age-related increase in sleep duration in AD patients (Figure 1b). We thus conclude that Aβ expression in flies models sleep-related changes in AD.

**Figure 2.**
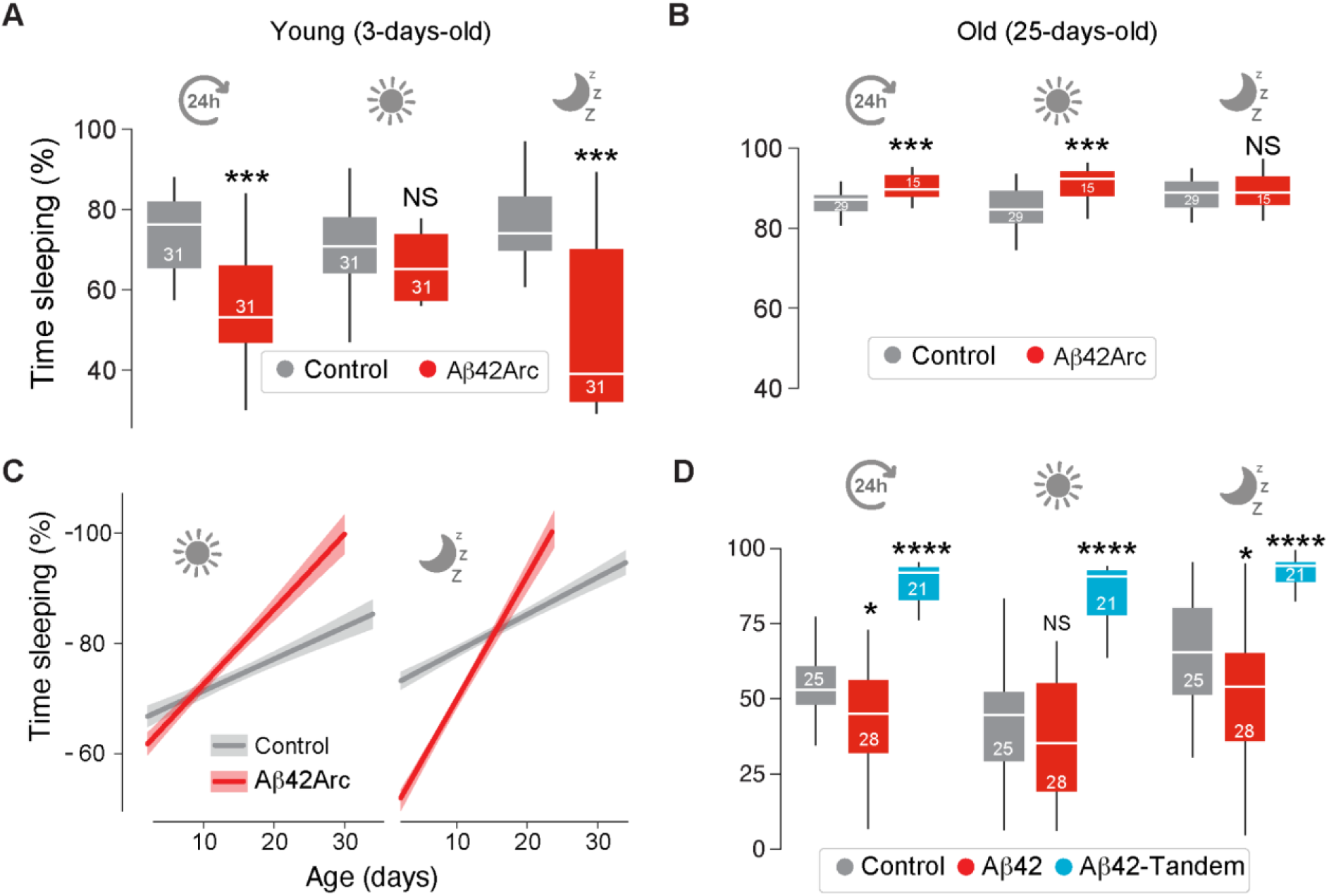
Distinct Aβ conformations lead to opposite outcomes with respect to sleep duration. **A**, Young flies (3 days old) expressing Aβ42Arc sleep less in the dark phase than do matched controls (means ± SDs; asterisks, two-tailed Student’s t test). **B**, Aged (25 days old) flies expressing Aβ42Arc sleep more than do matched controls (means ± SDs; asterisks, two-tailed Student’s t test). **C,** Age leads to increased sleep duration in flies expressing Aβ42Arc. **D,** Monomeric versus dimeric (tandem) Aβ forms cause divergent sleep outcomes in flies (means ± SDs; asterisks, two-tailed Student’s t test with Benjamini‒Hochberg correction). The expression of either monomeric or tandem Aβ42 was induced via the RU486 GeneSwitch system in 10-day-old flies. Sleep duration was recorded after 5 days of feeding the flies an RU486-supplemented diet. Genotypes: (A, B and C) *elav*Gal4; +; + (Control), *elav*Gal4; *UAS_Aβ42Arc/+*; + (Aβ42Arc) (D) *w*; +; *elav-GS-*Gal4/+ (Control), w; *UAS_Aβ42/+*; *elav-GS-*Gal4/+ (Aβ42), and *w*; *UAS_Aβ42-Tandem/+*; *elav-GS-*Gal4/+ (Aβ42-Tandem).

Next, we investigated the molecular mechanisms underlying the age-dependent changes in sleep duration caused by Aβ in *Drosophila*. In this context, older Aβ-expressing flies can sleep more because of increased aggregation of Aβ or an interaction between Aβ and ageing. We modelled our observations via the following equation: *Sleep duration ∼ Ageing_factors * β_1_ + Aβ_form * β_2_ + Ν*, where sleep duration in AD is determined by the effect (*β*) of a combination of ageing-related factors (*Ageing_factors*β_1_*) and the Aβ aggregation state (*Aβ_form * β_2_*), with some intrinsic biological noise (*Ν*).

To explore the effects of *Ageing_factors* and *Aβ_form*, we induced the expression of different forms of Aβ in adult flies aged 10 days via the GeneSwitch system. We observed that the expression of the monomeric Aβ42 peptide reduced sleep, whereas the expression of the dimeric Aβ42-Tandem peptide (Speretta et al., 2012) increased sleep duration (Figure 2d), suggesting that the *Aβ_form* is causally linked to Aβ-related sleep changes. Taken together, our results indicate that preclinical forms of AD are linked to reduced sleep, whereas increased AD progression increases sleep duration, possibly due to Aβ accumulation.

### Metabolomic signatures of the different forms of Aβ converge on NAD^+^

Expressing monomeric Aβ42 in flies decreases sleep duration without causing substantial neurodegeneration (Iijima et al., 2008), which mimics the preclinical phenotype in humans. Conversely, expressing an aggregate-prone version of Aβ42 (Aβ42rc) increases sleep duration and causes neurodegeneration (Yu et al., 2021a), which mimics a later stage of AD. We first investigated the metabolomic differences between flies expressing the two different forms of Aβ and controls, focusing on the metabolic changes that were uncoordinated between these flies (Figure 3a). We performed liquid chromatography-mass spectrometry (LC‒MS) on age- and sex-matched flies expressing the monomeric form (Aβ42) and compared them with those expressing Aβ42Arc, which is more prone to aggregation. We detected changes in 333 of the 454 detected metabolites (Figure 3a). We also found that metabolites linked to ROS levels (benzoate, γ-glutamylphenylalanine and γ-glutamyl-ε-lysine) as well as several metabolites linked to NAD^+^ metabolism were differentially abundant (Figure 3b). We previously showed that NAD^+^ levels are decreased in Aβ-expressing flies and that restoring NAD^+^ levels by inhibiting the NAD^+^-consuming protein Parp restored sleep defects (Yu et al., 2021a). A comparison of the metabolome of flies expressing these two forms of Aβ revealed a reversal in components of NAD^+^ metabolism. In flies expressing the monomeric form of Aβ, the levels of the NAD^+^ precursors nicotinamide riboside and nicotinamide mononucleotide decreased, whereas the level of NAD^+^ increased (Figure 3c, left). Conversely, in flies expressing the aggregate-prone form of Aβ, the levels of the NAD^+^ precursors nicotinamide riboside and nicotinamide mononucleotide increased, whereas the NAD^+^ level decreased (Figure 3c, right). Taken together, our comparison of flies expressing these two forms of Aβ points to an altered redox signature. γ-Glutamyl phenylalanine and γ-glutamyl-ε-lysine are associated with glutathione metabolism, where γ-glutamyl amino acids play crucial roles in the gamma‒glutamyl cycle (Wu et al., 2004). Lower NAD^+^ levels can also enhance ROS production by impairing the electron transport chain in mitochondria.

**Figure 3.**
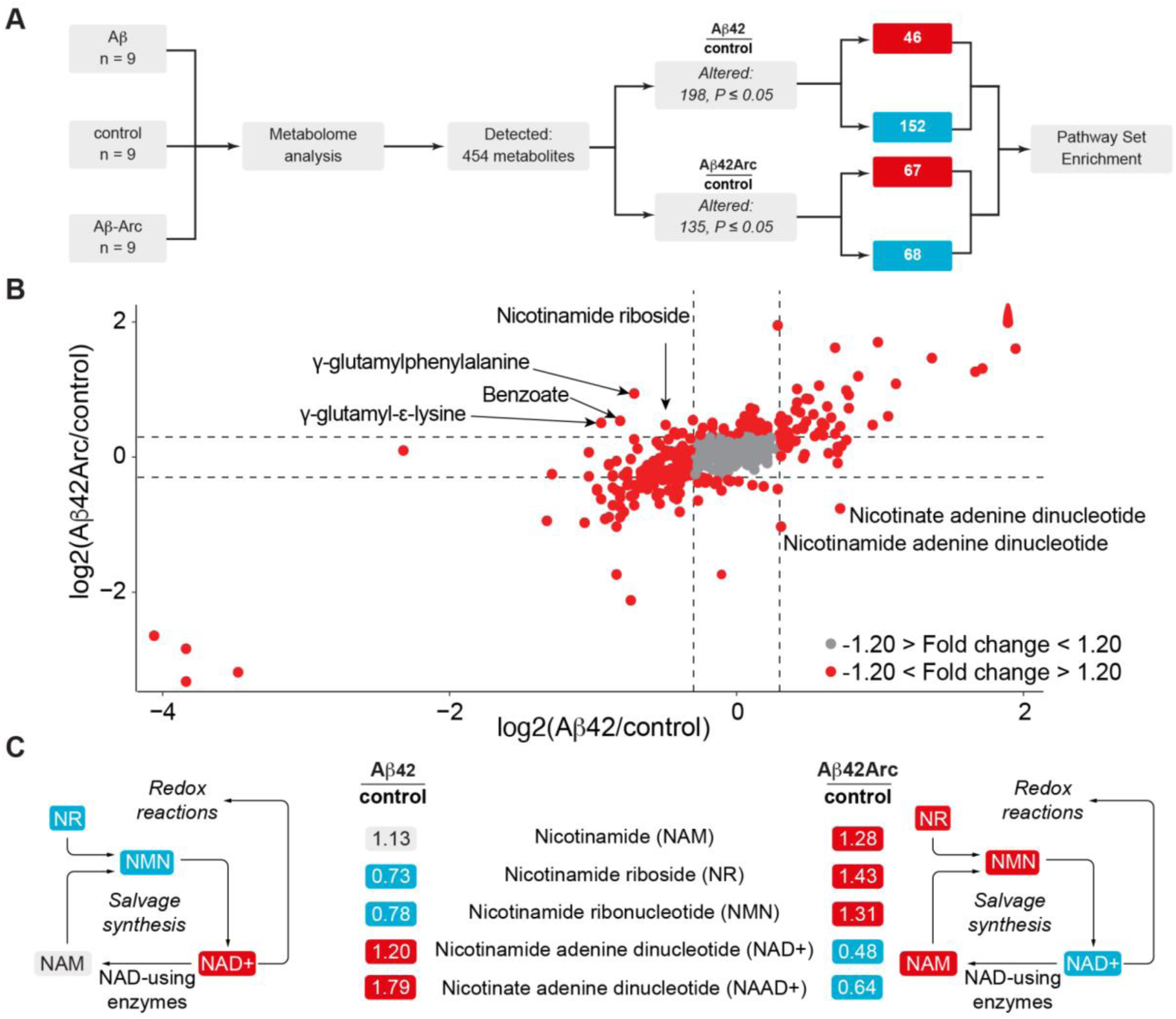
NAD^+^-linked redox metabolism is correlated with sleep changes in *Drosophila* models of AD. **A,** Workflow for the comparative analysis of metabolomic changes between control and either Aβ42- or Aβ42Arc-expressing flies. The fold changes of the detected metabolites were compared with those of the controls and analysed integratively. Significance was determined via Welch’s two-sample t test (n indicated for each sample set). **B,** Nicotinamide adenine dinucleotide metabolism and redox-related metabolites correlate with sleep changes in flies expressing the two different Aβ forms. Metabolites that changed above a 1.2-fold change are highlighted in red, and inconsistent differences between Aβ42- and Aβ42Arc-expressing flies are labelled. **C,** Schematic representation of the changes in NAD^+^ metabolism in Aβ42- and Aβ42Arc-expressing flies. Significantly increased and decreased metabolites are shown in red and blue, respectively. Metabolites that are not significantly changed are in grey. This figure is related to Supplementary Table 1. Genotypes: *elav*Gal4; +; + (Control), *elav*Gal4; *UAS_Aβ42Arc/+*; + (Aβ42Arc) and *elav*Gal4; *UAS_Aβ42/+*; + (Aβ42).

The altered metabolic signature in flies expressing two different forms of Aβ converges on ROS. Previous research has shown that ROS levels in the brain regulate sleep (Kempf et al., 2019). Therefore, we next measured the levels of ROS in the brains of flies expressing the two forms of Aβ employed in our study. Previous research has shown that higher ROS levels increase sleep duration, and in line with this finding (Kempf et al., 2019), we found that flies expressing Aβ42Arc, which have a longer sleep duration, have increased ROS levels in the brain (Figure 4a and b). We also detected a small, although not statistically significant, reduction in ROS in Aβ42-expressing flies compared with controls (Figure 4b). These results suggest that aggregate-prone Aβ42 increases sleep duration through increasing ROS levels. Consistently, we previously showed that the genetic suppression of ROS in Aβ42Arc-expressing flies restored the Aβ42Arc-related increase in sleep duration and increased lifespan (Yu et al., 2024). Overall, we conclude that aggregate-prone Aβ42Arc increases ROS, which increases sleep duration.

**Figure 4.**
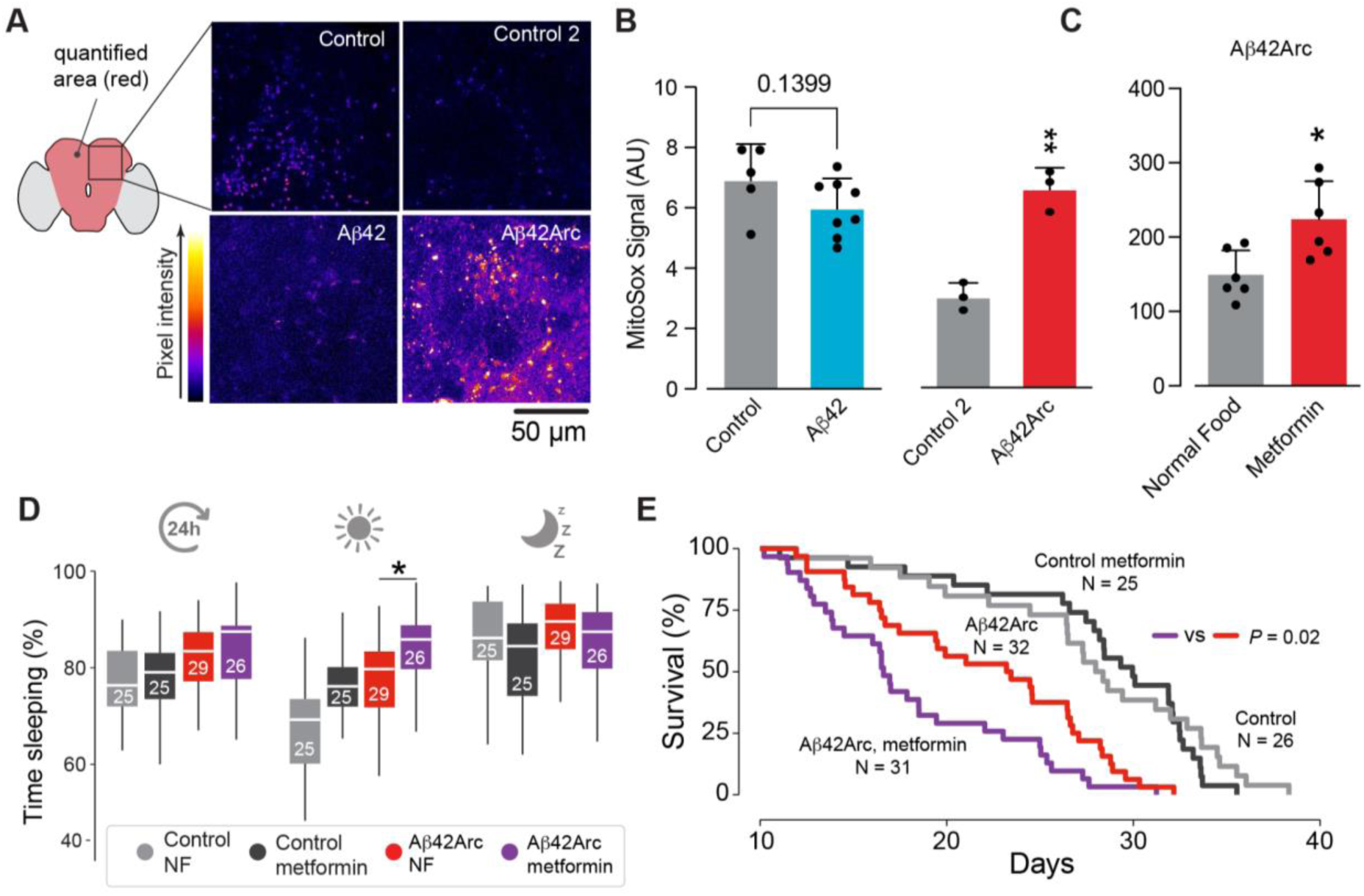
Metformin increases ROS and sleep and decreases lifespan in a *Drosophila* model of AD. **A** and **B,** Aβ42Arc-expressing flies have increased mitochondrial ROS in the brain. Representative confocal images (**A**) and quantitative analysis (**B**) of mitochondrial ROS in 10-day-old flies with the indicated genotypes (means ± SDs; asterisks, two-tailed Student’s t test). **C,** Aβ42Arc-expressing flies fed a metformin-containing diet presented increased ROS levels in the brain. Ten-day-old flies were maintained on a metformin-containing diet for 5 days before being assayed. **D**, A metformin-containing diet increased sleep duration during the light phase (middle) in both Aβ42Arc-expressing flies and controls compared with a normal diet (median ± IQR; asterisks, Kruskal‒Wallis rank sum test with Dunn’s test, adjusted for multiple comparisons via the Benjamini‒Hochberg method). **E**, A metformin-containing diet decreased the lifespan of Aβ42Arc-expressing flies (asterisks, log-rank test). For C and D, the flies were aged to 10 days post-eclosion in normal food and switched to a diet supplemented with 0.1 mM metformin for behavioural recording. Genotypes: *elavGal4; +; +* (control)*, elavGal4; +; UAS Aβ42Arc/+* (Aβ42Arc), and *elavGal4; UAS Aβ42/+; +* (Aβ42).

We next sought to further validate this conclusion via a clinically relevant paradigm. Metformin is a widely prescribed drug for type 2 diabetes with potential antiaging effects that has been used to treat AD (Imfeld et al., 2012; Khezri et al., 2022). However, metformin inhibits mitochondrial complex I at the ubiquinone-binding site (Bridges et al., 2023), which can increase ROS production. Expressing Aβ42Arc in flies also results in complex I impairment (Yu et al., 2024). We thus hypothesised that metformin exacerbates Aβ42Arc-related sleep pathologies. We fed 10-day-old Aβ42Arc-expressing flies metformin for 5 days and found a significant increase in ROS (Figure 4c) and sleep duration (Figure 4d). Feeding adult flies metformin also decreased their lifespan (Figure 4e). Taken together, these results indicate that higher ROS levels increase sleep duration changes in *Drosophila* models of AD.

### Altered protein signatures in flies expressing two different forms of Aβ

The expression of Aβ42Arc and Aβ42 in adult flies alters the NAD^+^ level, which plays a central role in cellular metabolism. To explore the protein changes linked to the altered NAD^+^ metabolism caused by Aβ42Arc and Aβ42, we used quantitative proteomics. We compared the proteomes of flies expressing these two forms of Aβ (Figure 5a) and found that 931 out of 5717 detected proteins were altered in both Aβ42Arc- and Aβ42-expressing flies compared with controls (Figure 5b). We next performed a gene ontology analysis based on the molecular functions of the differentially abundant proteins. We found significant enrichment in several protein networks, including networks of enzymes that use NAD^+^ or NADP^+^ as coenzymes (Figure 5c), such as malate dehydrogenase 1, glucose-6-phosphate 1-dehydrogenase (with the gene name Zw) and hyperkinetic.

**Figure 5.**
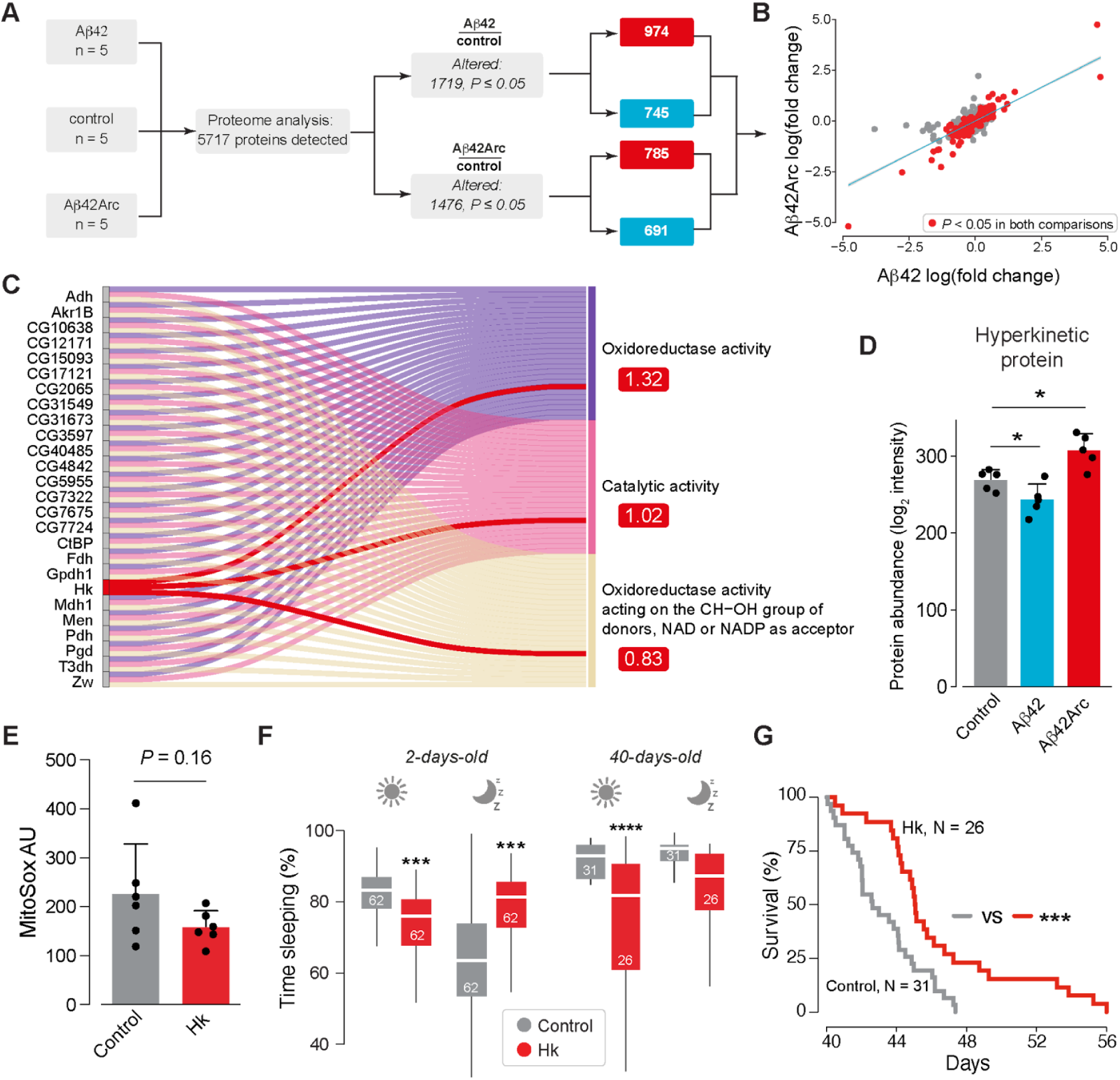
The redox sensor Hyperkinetic is linked to Aβ and sleep duration in flies. **A,** Workflow for the comparative analysis of proteomic changes between control and either Aβ42- or Aβ42Arc-expressing flies. Five biological replicates containing approximately 20 fly heads each were homogenised and analysed by mass spectrometry. Male flies aged 10 days post-eclosion were used. **B,** Proteome-wide comparison of differentially expressed proteins associated with Aβ42 on the x-axis and Aβ42Arc on the y-axis to those associated with the controls. Proteins with a fold change that was statistically significant in both comparisons (Aβ42 vs. control and Aβ42Arc vs. controls) are labelled in red, whereas others are labelled in grey. **C,** Pathway analysis of the differentially abundant proteins in both conditions at nominal significance (labelled in red in **B**). The analysis was performed based on molecular function enrichment, and only the top 3 pathways are shown. Significant pathways are on the right side, and proteins linked to the enriched pathways are shown. Functional enrichment was performed on the proteins matched to the STRING database (915 out of 931). The values in red boxes are the signal values from STRING, which is the weighted harmonic mean between the observed/expected ratio and -log(FDR). The connections between Hk and its associated pathways are highlighted in red. **D,** Hk was increased in Aβ42Arc-expressing flies but decreased in Aβ42-expressing flies. Protein levels were measured via mass spectrometry (means ± SDs; asterisks, two-tailed Student’s t test, adjusted for multiple comparisons via the Benjamini‒Hochberg method). **E,** ROS levels are not altered in Hk-overexpressing transgenic flies (means ± SDs; asterisks, two-tailed Student’s t test). Ten-day-old flies were used. **F,** Hk overexpression in young flies decreases sleep during the light phase and increases sleep during the dark phase. Hk overexpression decreases sleep during the light phase in old flies (means ± SDs; asterisks, two-tailed Student’s t test). **G**, Hk overexpression increases lifespan (means ± SDs; asterisks, log-rank test). The lifespan was recorded for flies aged 40 days after eclosion. This figure is related to Supplementary Table 2. Genotypes: *elavGal4; +; +* (Control) and *elavGal4; UAS Hk/+; +* (Hk).

The NAD^+^ level was increased in flies expressing Aβ42 and decreased in flies expressing the aggregate-prone form of Aβ42. The overexpression of Aβ42 decreased sleep duration, whereas the overexpression of Aβ42Arc increased sleep duration (Figure 5d). We therefore looked for protein abundances in our network that were reversed under both conditions. We found that the levels of hyperkinetic (Hk), which previous studies have shown to integrate redox and metabolic changes with behaviour (Bushey et al., 2007; Kempf et al., 2019; Pimentel et al., 2016), matched this condition. Compared with the control flies, the protein level of Hk was lower in the Aβ42-expressing flies and higher in the Aβ42Arc-expressing flies (Figure 5d).

Hk loss of function results in shorter sleep cycles and cognitive impairment (Bushey et al., 2007). We found that the pan-neuronal overexpression of Hk in wild-type flies was linked to a small, nonstatistically significant decrease in ROS (Figure 5e). Hk overexpression led to a decrease in sleep duration during the light phase and an increase in sleep duration during the dark phase (Figure 5f, left). In older flies, Hk overexpression significantly increased wakefulness during the light phase (Figure 5f, right) and improved lifespan (Figure 5g). We next tested whether Hk overexpression could protect against the toxic effects of Aβ42Arc expression. The overexpression of Hk in Aβ42Arc-expressing flies reduced the levels of ROS in the brain (Figure 6a), increased wakefulness (Figure 6b), and was neuroprotective (Figure 6c and 6d). We conclude that Hk overexpression is neuroprotective in a fly model of AD.

**Figure 6.**
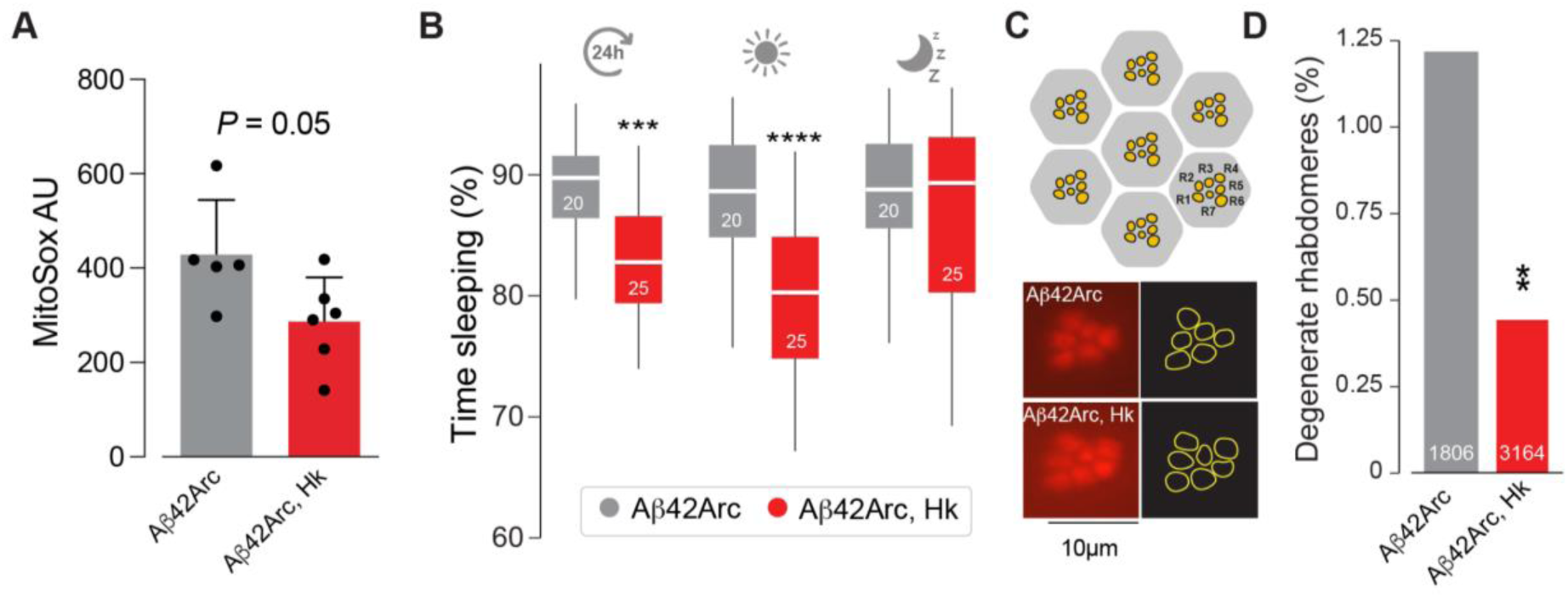
Increasing the expression of *Hyperkinetic* increases daytime activity and delays neurodegeneration in a *Drosophila* model of AD. **A**, Hk overexpression in Aβ42Arc-expressing flies reduces the levels of mitochondrial ROS in the brain (mean ± SD; asterisks, two-tailed Student’s t test). **B,** Hk overexpression in Aβ42Arc-expressing flies increased wakefulness (mean ± SD; asterisks, two-tailed Student’s t test). Sleep duration was recorded for 5 days in 10-day-old flies. **C**, An illustration of the typical layout of visible photoreceptors (red, R1–R7) at the surface of adult *Drosophila* ommatidia. A healthy ommatidium has 7 visible rhabdomeres, while neurodegeneration usually leads to a lower number of visible rhabdomeres. **D**, Hk overexpression rescues the neurodegeneration of photoreceptor cells in Aβ42Arc-expressing flies (asterisks, two-tailed chi-square test, 95% confidence interval). Genotypes: *elavGal4; +; UASAβ42Arc/+* (Aβ42Arc) and *elavGal4; UAS Hk/+; UAS Aβ42Arc/+* (Aβ42Arc, Hk). All flies were 10 days post-eclosion when tested.

### *KCNAB1* and *KCNAB2* expression at single-cell resolution follows AD progression

Fly *Hk* has 2 orthologues in humans (*KCNAB1* and *KCNAB2*), which encode voltage-gated potassium channel subunits β with conserved structures and functions (Gulbis et al., 1999). *KCNAB2* haploinsufficiency is associated with neuronal dysfunction in patients (Heilstedt et al., 2001). We observed that protein levels of Hk are decreased in flies expressing monomeric Aβ, whereas they are increased in flies expressing aggregation-prone Aβ (Figure 5d). On the basis of the assumption that KCNAB protein levels are positively correlated with their expression levels, we hypothesised that the *KCNAB* levels might follow the trend of Hk in flies expressing different forms of Aβ42. Specifically, the levels of *KCNAB* might be decreased in neurons from patients with low levels of AD pathology and increased in patients with a high AD burden. Hk levels are elevated in inhibitory neurons from the *Drosophila* dorsal fan-shaped body (Ni et al., 2019); therefore, we analysed the expression of *KCNAB1* and *KCNAB2* at the single-cell level in human inhibitory neurons. We first gathered and compared data from patients with clinically confirmed AD and classified as Braak stage III or above with data from non-AD individuals (Otero-Garcia et al., 2022). Using a combination of univariate and multivariate analyses, we found that the expression levels of *KCNAB1* and *KCNAB2* were greater in patients with AD than in individuals without an AD diagnosis (Figure 7a–d). We next modelled a low level of AD pathology on the basis of Braak staging consistent with the preclinical stage of AD, with no clinically significant symptoms described in Figure 1. Individuals classified as Braak stage I have evidence of neurofibrillary tangles only in the entorhinal region of the brain (Braak et al., 2006) but do not show any memory impairment (Therriault et al., 2022). To model the at-risk and preclinical stages of AD, we compared gene expression data at the single-cell level of inhibitory neurons from post-mortem brains from individuals with Braak stage I with those from Braak stage 0. We detected a significant decrease in *KCNAB1* and *KCNAB2* expression levels in both the univariate and multivariate models (Figure 7e–h), which aligns with our results in flies expressing monomeric Aβ (Figure 5d). Overall, we conclude that at the preclinical stage of AD, *KCNAB1* and *KCNAB2* expression levels are decreased. As the disease progresses, *KCNAB1* and *KCNAB2* expression increases, establishing a parallel relationship between the expression levels of *KCNAB1/2* in the human brain and Hk in our *Drosophila* model of AD.

**Figure 7.**
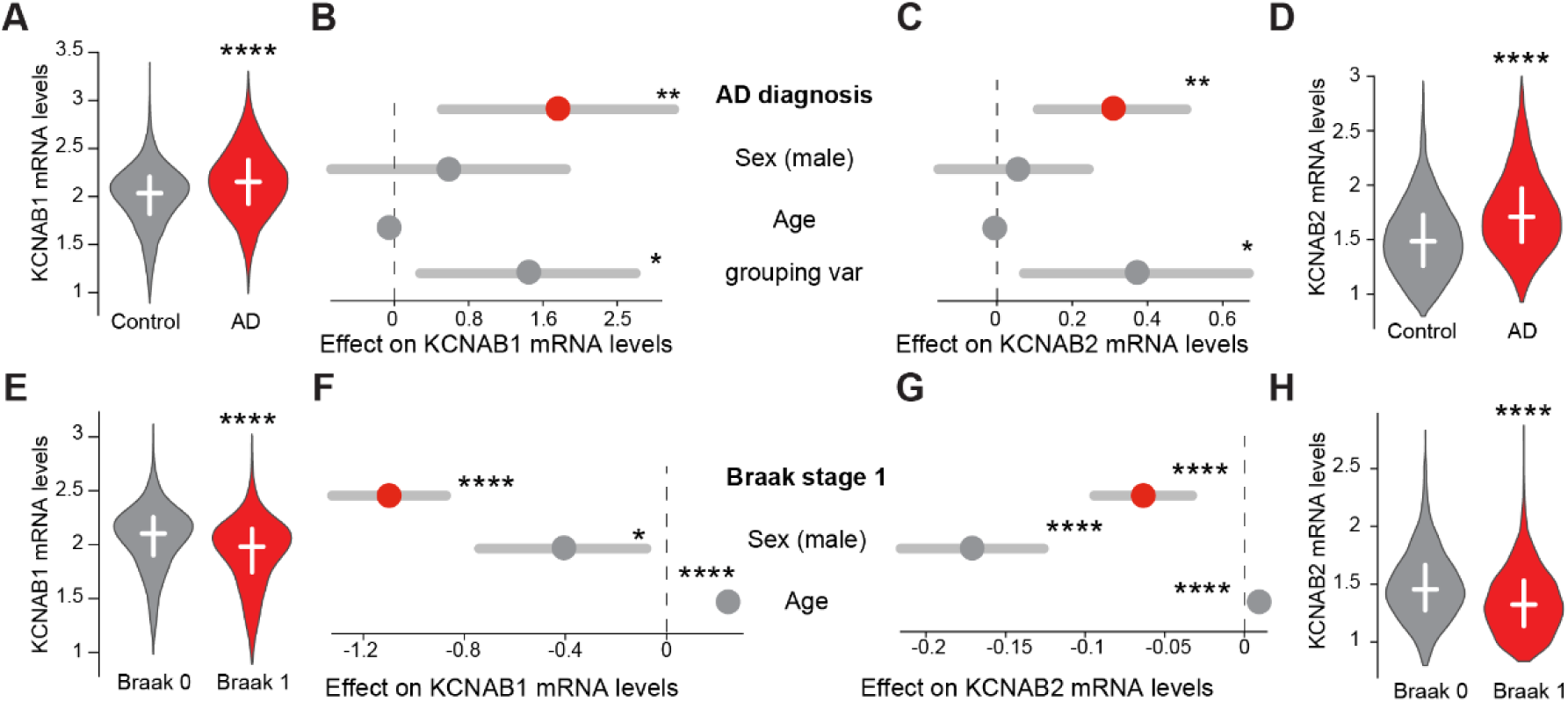
*KCNAB1* and *KCNAB2* expression levels are associated with AD pathology in inhibitory neurons. **A‒D**, Inhibitory neurons from AD patients have increased expression of *KCNAB1* and *KCNAB2*. **A, B**, *KCNAB1* expression was increased in single-cell RNA sequencing data from AD patients (**A,** median ± IQR; asterisks, two-tailed unpaired Mann–Whitney test) and remained significant when covariates were accounted for (**B,** asterisks, linear mixed model). **C, D**, *KCNAB2* expression was increased in single-cell RNA sequencing data from AD patients according to both adjusted and univariate models (**C,** asterisks, linear mixed model; **D,** median ± IQR; asterisks, two-tailed unpaired Mann‒Whitney test). **E‒H,** *KCNAB1* and *KCNAB2* expression levels are lower in the neurons of donors with mild AD pathology, modelled as Braak stage 1, than in those of donors categorised as Braak stage 0 (no AD pathology). **E, F,** KCNAB1 expression was decreased in single-cell RNA sequencing data from Braak stage 1 donors (**E,** median ± IQR; asterisks, two-tailed unpaired Mann‒Whitney test) and remained significant when covariates were accounted for (**F,** asterisks, linear regression model). **C, D**, KCNAB2 expression was also lower in single-cell RNA sequencing data from Braak stage 1 patients than in data from Braak stage 0 patients according to both adjusted and univariate models (**C,** asterisks, linear mixed model; **D,** median ± IQR; asterisks, two-tailed unpaired Mann‒Whitney test). For Panels **B, C, F,** and **G**, the error bars in grey represent the 95% CIs of the estimated effect (dot). The investigated variable is in red.

### Mendelian randomisation indicates that the brain expression of *KCNAB2* decreases AD risk

Given the parallel established in the previous section and the protective effect of the upregulation of Hk in a fly model of AD (Figure 6), we hypothesised that increased *KCNAB1* and *KCNAB2* expression could also be protective in humans. Mendelian randomisation (MR) uses genetic variations (single nucleotide polymorphisms, SNPs) linked to an exposure to examine the causal effect of that exposure on an outcome (Sanderson et al., 2022). By using SNPs associated with *KCNAB* expression in the brain, we used MR to determine whether higher predicted *KCNAB* levels are linked to AD risk (Figure 8a). Compared with observational studies, MR studies are less prone to bias and confounding factors and can provide evidence regarding causality. We found that SNPs linked to higher *KCNAB2* protein levels in the dorsolateral prefrontal cortex (Wingo et al., 2021) are also significantly linked to a decreased risk of developing AD (Figure 8b), indicating that higher *KCNAB2* protein expression protects against AD risk. Given our previous analysis of *KCNAB2* mRNA levels in inhibitory neurons, we next modelled the effect of *KCNAB2* expression specifically in inhibitory neurons (Bryois et al., 2022). We found that increased *KCNAB2* expression was also linked to decreased AD risk (Figure 8c), supporting our previous results. Overall, we conclude that increased neuronal expression of *KCNAB2* decreases AD risk in humans, establishing a parallel between the expression of KCNAB2 in AD brains and the effects of Hk upregulation in *Drosophila* models of AD associated with Aβ toxicity.

**Figure 8.**
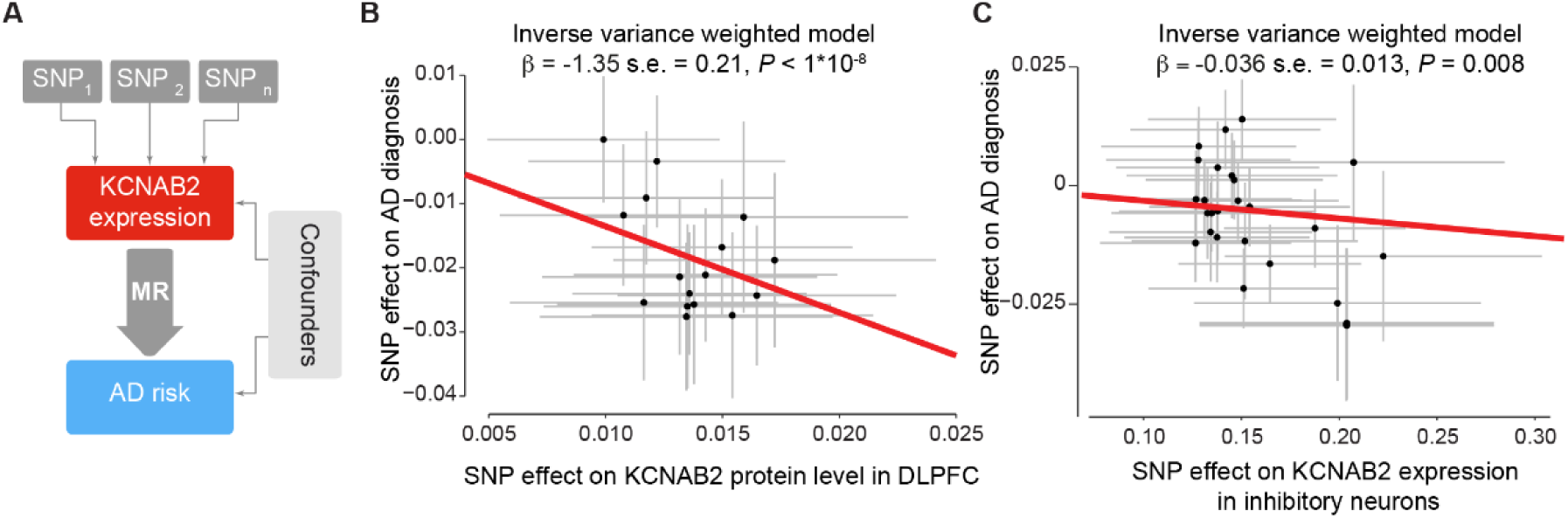
Higher KCNAB2 expression decreases AD risk. **A**, MR uses SNPs to infer the causal effect of gene expression on AD risk. **B,** Increased KCNAB2 protein levels in the dorsolateral prefrontal cortex (DLPFC) decrease AD risk. **C**, Increased *KCNAB2* mRNA levels in inhibitory neurons decrease AD risk.

## Discussion

Sleep problems are often thought to cause or worsen AD. Here, we showed that AD pathology might also drive gradual changes in sleep duration, possibly through the modulation of redox metabolism in flies. We previously established that enhancing NAD^+^ levels and reducing oxidative stress restored sleep pathology in *Drosophila* models of AD (Yu et al., 2024, 2021a), but the underlying mechanisms were unclear. On the basis of these findings, we show that NAD^+^-linked redox metabolism and the homeostatic sleep switch are associated with sleep disturbances in AD patients. These findings suggest that suppressing ROS by improving mitochondrial function could alleviate sleep disturbances in AD patients.

We modelled a preclinical stage of AD via a PRS in the UK Biobank analysis (McKhann et al., 2011). *APOE* alleles were the main contributors to the PRS used in our study. The *APOE* gene, which encodes apolipoprotein E, is involved in lipid metabolism, synaptic function, neuroinflammation and Aβ accumulation (reviewed in Kim et al., 2009). The ε4 allele of *APOE* is the major genetic risk factor for AD and is linked to a greater Aβ load in the brain (Kanekiyo et al., 2014). We showed that individuals with a high PRS have decreased sleep duration (Figure 1), mimicking findings in flies expressing monomeric Aβ (Figure 2). However, our longitudinal analysis of sleep duration in flies spanned 40 days, whereas our UK Biobank analysis was cross-sectional. With additional longitudinal data collected from the UK Biobank participants, future work could determine whether patients with an AD diagnosis reported shorter sleep durations prior to clinical symptoms such as cognitive decline. This, together with longitudinal imaging of Aβ in the brains of AD patients through methods such as positron emission tomography (Lopes Alves et al., 2020) and a correlation with the sleep duration of individuals from their period before and after AD diagnosis, would also contribute to confirming our findings.

To confirm our results on the dynamic changes in sleep duration across AD pathologies (Figure 1), we established a parallel between our fly models of Aβ and found an age-related increase in sleep duration (Figure 2). As flies age, age-related changes could occur that moderate the effects of Aβ. Alternatively, with increasing age, monomeric Aβ could aggregate and cause different effects. By comparing the monomeric and dimeric forms of Aβ, we showed that different Aβ forms moderate sleep duration in flies (Figure 2d). We conducted exploratory analyses by comparing the proteome and metabolome of monomeric and aggregate-prone forms of Aβ. The use of Aβ42Arc, rather than Aβ42-Tandem, is primarily justified by the nearly identical amino acid sequences of Aβ42Arc and the monomer, differing only by a single amino acid substitution that promotes aggregation. Given their identical size and shared genomic locus (Moloney et al., 2010), the Aβ42 and Aβ42Arc trans genes facilitate controlled comparative analysis. The expression of Aβ42-Tandem causes substantial lethality to flies (Luheshi et al., 2007), which limits the generation of samples for proteomic and metabolomic analyses. Additionally, employing Aβ42-Tandem-expressing flies requires the use of the GeneSwitch system, which relies on keeping flies in a diet supplemented with a drug, RU486, to induce expression of the trans gene. This method introduces additional noise in downstream analyses, as flies can stochastically eat different amounts of food, thereby increasing variability, which is detrimental to high-precision omics analyses. Feeding flies RU486 also alters the expression of mitochondria-related genes (Robles-Murguia et al., 2019), which can exacerbate the effect of Aβ. These considerations led us to use the GAL4-UAS system (Brand and Perrimon, 1993) to induce the expression of monomeric Aβ42 and aggregate-prone Aβ42Arc in our experiments.

The metabolomics data suggested variations in NAD^+^ metabolism and redox potential between flies expressing either Aβ42 or Aβ42Arc. Specifically, Aβ42Arc-expressing flies presented increased mitochondrial ROS, whereas Aβ42-expressing flies presented no significant changes in ROS levels. The observed minimal decrease in mitochondrial ROS in Aβ42-expressing flies may reflect limitations in the sensitivity of the mitoSOX ROS detection method. Mass spectrometry could be employed in future studies to assess glutathione metabolism more accurately as a proxy for the redox state.

Here, we observed an increase in ROS in the brain by feeding Aβ-expressing flies metformin, a complex I inhibitor (Figure 4), which led to an increase in sleep duration, indicating a causal role of ROS signalling in sleep duration (Hill et al., 2018). Although these findings indicate a potential toxic effect of metformin in advanced AD, its clinical significance remains uncertain because of the multifaceted effects of metformin on multiple organs (Foretz et al., 2023; Khezri et al., 2022). Future clinical research on the effects of metformin on AD should closely monitor sleep disturbances in patients in light of the potential side effects reported.

Proteomic analysis revealed a potential association between Hk, an NADP-dependent sleep regulator, and AD pathogenesis. We did not find suitable antibodies to confirm the changes in Hk levels detected by proteomics. Instead, we sought to test this finding in a clinically relevant way, using human post-mortem neurons, on the basis of the assumption that mRNA and protein levels are correlated. Using inhibitory neurons from a small number of donors, we showed that neurons from donors with a low level of AD pathology (Braak stage I) have lower levels of both *KCNAB1* and *KCNAB2* expression, whereas those from AD donors (Braak stage IV) have higher expression levels. Population-scale single-nucleus RNA sequencing datasets from AD patients are now available (Mathys et al., 2023); therefore, future work can analyse *KCNAB* expression in a larger dataset across disease severity, cell types and brain regions. Genetic analyses via MR revealed that increased KCNAB2 expression in the brain at both the mRNA and protein levels decreased AD risk (Figure 8), corroborating our results in flies (Figure 6). We did not analyse the effect on *KCNAB1* protein or mRNA expression because of the lack of SNPs associated with their expression. Larger single-cell transcriptomics datasets will enable more SNPs associated with *KCNAB* expression to be identified and help validate our findings.

Research in flies has shown that homeostatic sleep switches Hk levels, connecting animal neuronal function and redox to behaviour (Bushey et al., 2007; Kempf et al., 2019; Pimentel et al., 2016). Our findings indicate that increased Hk levels play an adaptive and protective role in AD by increasing sleep duration. We found that Hk protein abundance was increased in the AD model with increased ROS levels (Figure 5), which aligns with previous research showing that Hk expression is increased in subcellular locations with increased ROS (Ueda and Wu, 2008). Hk overexpression in young flies caused an increase in wakefulness during the light phase and an increase in sleep during the dark phase (Figure 5f). In flies, ROS production is increased during the light phase (Krishnan et al., 2008). Therefore, the overexpression of Hk in flies might cause a decrease in ROS, thus leading to increased wakefulness. During the dark phase, when ROS levels are lower, higher levels of Hk might increase sleep-promoting slow inactivation of A-type currents (Kempf et al., 2019). In older flies and flies expressing Aβ, the overexpression of Hk increased wakefulness during the light phase (Figure 5f). We found that Aβ42Arc-expressing flies have increased ROS levels in the brain, and increased ROS levels are also found in the brains of aged flies (Gomes et al., 2023). Therefore, with elevated ROS, Hk overexpression might decrease ROS and consequently increase wakefulness. Future research could explore how the redox function of Hk regulates neuronal activity and sleep in fly models of AD. For example, this could be done by comparing the overexpression of Hk to a version containing a point mutation (K289M) that abolishes its oxidoreductase activity (Fogle et al., 2015).

Although our work focused on the relationship between mitochondria and Hk in the context of sleep and AD, the inhibition of mitochondrial respiration complexes can also affect the circadian component of sleep. TIMELESS is a circadian clock protein that contributes to maintaining and regulating circadian rhythms (Sehgal et al., 1995). Mitochondrial complex I is impaired in models of AD (Yu et al., 2024), and the inhibition of complex I facilitates the degradation of TIMELESS (Zheng et al., 2024), which can disrupt circadian rhythmicity. A TIMELESS-mediated circadian disruption might explain the disrupted circadian rhythmicity in AD (Musiek et al., 2015).

Glial cells can also drive AD pathogenesis through neuroinflammation (Heneka et al., 2024). In flies, mitochondrial oxidation in glial cells also regulates sleep patterns (Haynes et al., 2024). Future work can investigate the impact of Aβ aggregation forms on the cellular and molecular dynamics between glia and neurons and the behavioural effects on sleep duration.

Our study does not explain the observed alterations in Hk levels between flies expressing two different forms of Aβ. One possibility is that different Aβ forms inhibit different proteins, causing cellular stress and thereby affecting Hk levels. To test this hypothesis, future work could employ thermal proteomic profiling to identify the binding partners of different forms of Aβ (Mateus et al., 2020). Different tau structures are also present in AD patients and may lead to diverse pathologies (Dujardin et al., 2020; Shi et al., 2021). The aggregated forms of TAR DNA-binding protein 43 kDa, which can cause amyotrophic lateral sclerosis, are also diverse (Arseni et al., 2023, 2022), suggesting that the distinct aggregated forms of toxic proteins or peptides can lead to different cellular phenotypes. Therefore, understanding the toxic consequences of different conformations of these aggregate-prone molecules is necessary to increase our understanding of progressive neurodegenerative diseases.

## Author Contributions

Y.Y. initiated the project. Y.Y., S.H.Y.L. and L.M.M. designed the study, coordinated the experiments and provided conceptual input for the paper. Y.Y. performed the experiments. I.C. contributed to the generation of the metabolomics data. Y.Y. performed the computational analyses with help from B.W.Z.T. and L.Z. Y.Y. and L.M.M. wrote the manuscript with help from G.F. All the authors read and approved the final manuscript.

## Acknowledgements

Mass spectrometry analysis was performed at the Proteomics Facility of the Medical Research Council Toxicology Unit University of Cambridge, Cambridge, UK. The authors would like to thank Dr. Catarina Franco and Dr. Bini Ramachandran for their help with the preparation and data analysis of the proteomics samples. This work was funded by the UK Medical Research Council intramural project MC_UU_00025/3 (no. RG94521) to L.M.M. The funders had no role in the study design, data collection and analysis, decision to publish or preparation of the manuscript.

## Methods

### Genetics and *Drosophila* strains

Fly stocks and crosses were maintained on standard cornmeal agar media at 25 °C. The strains used were *elavGAL4*, *w; UAS Aβ42ARC; +* and *w; 51D; +* (described in Yu et al., 2021), *w; UAS Aβ42-Tandem; +*, *w; UAS Aβ42;+*, *w; +; UAS Aβ42ARC and w; +; 86F*, (kind gifts from D. Crowther, R&D Neuroscience, Innovative Medicines and Early Development Biotech Unit, AstraZeneca, Cambridge, UK), *w*; +; *elav-GS-*Gal4/+ (kind gift from A. Whitworth) and *w; UAS_Hyperkinetic; +* was obtained from Bloomington Stock Center: 86270 (Fogle et al., 2015; Kempf et al., 2019).

### Metabolic profiling

The results of the analysis of the global metabolomics of Aβ42Arc- and Aβ42-expressing flies compared with controls were obtained from the Metabolon Platform (Metabolon Inc., NC, USA). All female flies aged 15 days after eclosion were used. The data from Aβ42Arc-expressing flies were previously published (Yu et al., 2021a), the data from Aβ42-expressing flies were acquired at the same time via the same method, and the same statistical workflow was used.

### Drug treatments

Metformin (PHR1084, Sigma Aldrich) was freshly prepared as a 1 M stock in water and dissolved in fly food at a final concentration of 0.1 mM. This concentration was chosen on the basis of the work of Slack et al., 2012. Flies from each genotype were randomly assigned to normal food or supplemented food. To induce the expression of different Aβ forms, we used the GeneSwitch system. We aged the flies to 10 days on standard cornmeal agar media at 25 °C. To induce expression, we fed flies the drug RU486 (Abcam, ab120356) in standard fly food at a final concentration of 500 µM.

### Locomotor assays and lifespan analysis

The analyses of sleep and lifespan in flies were performed as described previously (Yu et al., 2021a). Adult male flies were individually loaded in glass tubes containing the same food used for rearing. The flies were grown and analysed under a light/dark cycle of 12 h/12 h at 25 °C. The total number of recorded midline crossings per minute was recorded via the Drosophila Activity Monitoring System (Trikinetics, Waltham, MA), and the data were analysed via Rethomics (Geissmann et al., 2019). The analysis started at the first ZT0 to allow acclimation. The duration of sleep was calculated for the first 5 days, and the data of the flies that died were discarded. Sleep was defined as 5 min of inactivity. The data for lifespan analysis are presented as Kaplan–Meier survival distributions. We provide the full analysis script and the raw data in our GitHub repository (https://yizhouyu.com/abeta_sleep).

### Microscopy-based assessment of mitochondrial reactive oxygen species

Measurements of ROS in fly brains were performed as previously described (Travaglio et al., 2023). The brains of 10-day-old male flies were dissected in cold PBS and incubated with 5 μM MitoSOX Red mitochondrial superoxide indicator (M36008, Molecular Probes) for 30 min. After incubation, the brains were washed with PBS for 5 min and immediately imaged on a Zeiss LSM880 confocal microscope. Stacks that were 89 μm thick were acquired. The maximal intensity projections of the MitoSOX signal in the fly midbrain were quantified via ImageJ.

### Pseudopupil analysis

The heads of the flies were directly fixed on standard microscope slides via quick-dry transparent nail varnish as previously described (Yu et al., 2021a). A Zeiss Axioplan 2 microscope (Zeiss BMT Bayerl Messtechnik GmbH, Germany) equipped with a 63x oil immersion objective was used to visualise the ommatidia. Approximately 5 flies per condition were examined to obtain approximately 1000 rhabdomeres. The percentage of abnormal rhabdomeres was calculated as the number of degenerate rhabdomeres over the total number of rhabdomeres: (A*1 + B*2 + C*3)/N, where A = the number of ommatidia with 6 rhabdomeres, B = the number of ommatidia with 5 rhabdomeres, C = the number of ommatidia with 4 rhabdomeres and N = the total number of ommatidia counted. Statistical significance was determined via a chi-square test.

### Proteome profiling

Proteomic analysis was performed as previously described (Fedele et al., 2022; Stefanatos et al., 2023). Briefly, 10 heads from 10-day-old adult males from each condition were homogenised in 450 µL of 100 mM triethylammonium bicarbonate (TEAB) for 5 cycles of 20 s at 5500 rpm with 10 s intervals at 4 °C. Next, 1% RapiGest buffer and a further 450 µL of 100 mM TEAB buffer were added to the samples, followed by sonication and centrifugation at 2000 × g for 5 min at 4 °C. A total of 850 µL of the supernatant was collected for further processing. Fifty micrograms of protein extract was digested in 50 µL of 100 mM TEAB. The samples were denatured at 80 °C for 10 min. A total of 2.8 μl of 72 mM DTT was added, and the samples were heated at 60 °C for 10 min. A total of 2.7 μl of 266 mM iodoacetamide was added, and the samples were incubated at room temperature for 30 min. The DTT concentration was increased to 7 mM to quench alkylation. The samples were digested with 1 µg of trypsin (5 µL of 0.2 µg/µL stock) overnight at 37 °C. Digests were quantified via the peptide quantification assay from the Pierce™ Quantitative Colorimetric Peptide Assay (Cat no: 23275) according to the manufacturer’s instructions. 2xTMT11 with pooled reference standards was used to multiplex the samples. LC‒MS/MS was performed via synchronous precursor selection (SPS MS3) with MS3-based quantification triggered by Drosophila MS2-identified peptide real-time search. The injected samples were analysed via an Ultimate 3000 RSLC™ nanosystem (Thermo Fisher, Hemel Hempstead) coupled to an Orbitrap Eclipse™ mass spectrometer (Thermo Fisher). Data were acquired via three FAIMS CVs (−45 V, −60 V and −75 V), and each FAIMS experiment had a maximum cycle time of 2 s. For each FAIMS experiment, the data-dependent SPSMS3 RTS program used for data acquisition consisted of a 120,000-resolution full-scan MS scan (AGC set to 50% (2e5 ions) with a maximum fill time of 30 ms) using a mass range of 415–1500 m/z. The raw data were imported and processed in Proteome Discoverer v2.5 (Thermo Fisher). The raw files were subjected to a database search via Proteome Discoverer with Sequest HT against the UniProt UP000000803 Drosophila database (containing 21135 seq. accessed on 2022/08/04). Common contaminant proteins (human keratins, BSA and porcine trypsin) were added to the database. Statistical analyses for fold change and significance were performed as previously described (Popovic et al., 2021). For functional enrichment analysis, we selected proteins altered in both conditions compared to control at nominal significance (*P* < 0.05 in both Aβ42 and Aβ42Arc compared to controls). We used this list to query the STRING database. The analysis source code and quality checks are available at GitHub (https://yizhouyu.com/abeta_sleep).

### UK Biobank analysis

The UK Biobank comprises health data from over 500,000 community volunteers based in England, Scotland and Wales. The geographical regions, recruitment and other characteristics have been previously described (Bycroft et al., 2018). Informed consent was obtained from all the subjects. UK Biobank ethical approval was granted by the North West Multi-Centre Research Ethics Committee. The current analysis was approved under UK Biobank application #60124.

For the analysis of the relationship between sleep duration and AD, we collected similar phenotypic data as in Yu et al. 2021, including data on sleepiness, polygenic risk for AD, age, waist‒hip ratio, sex, ethnicity, education level and social deprivation (Townsend deprivation index). Multivariate linear regressions were performed as previously described via R version 4.0.0 (Leal et al., 2021; Travaglio et al., 2021; Yu et al., 2021b). The polygenic risk score used to model AD risk was previously generated (Yu et al., 2021a). The analysis source code, detailed quality checks and all supplementary material are available at GitHub (https://yizhouyu.com/abeta_sleep).

### Analysis of single-cell RNA sequencing data

For the analysis of gene expression in single-cell RNA sequencing data, processed sequence reads (Otero-Garcia et al., 2022) were obtained and analysed via scanpy (Wolf et al., 2018). Preprocessing was performed by removing genes in fewer than 3 cells and cells with fewer than 200 genes. The mitochondrial gene expression cut-off was 2.52%, and cells above this cut-off were removed. Gene expression was normalised and ln(x+1) transformed. Cells with no KCNAB expression were removed.

Next, we used linear regression on single neurons to query the associations between the expression levels of *KCNAB* and their AD status or Braak stages, accounting for age and sex. For the analysis of patients with clinically diagnosed AD compared with controls, we used linear mixed models, which account for interindividual differences in gene expression and included age and sex as covariates. For the analysis of preclinical AD, we compared patients with Braak stage I disease with controls. We used linear regressions due to the low number of people with a Braak stage I label and included age and sex as covariates.

### Mendelian randomisation

For the brain-specific Mendelian randomisation analysis, we incorporated the exposure data of protein quantitative trait loci (pQTL) from the dorsolateral prefrontal cortex (Robins et al., 2021). We only considered statistically significant pQTL (*P* < 0.05) linked to *KCNAB2* expression in the dorsolateral prefrontal cortex. We did not find enough pQTL linked to *KCNAB1* expression.

For the cell-type-specific Mendelian randomisation analysis, we incorporated the exposure data of cell-specific expression quantitative trait loci (eQTL) (Bryois et al., 2022). Similarly, we considered only statistically significant eQTL (*P* < 0.05) linked to *KCNAB2* expression in inhibitory neurons. We combined these exposure data with the outcome data from the largest GWAS for AD (Bellenguez et al., 2022). We performed clumping to remove correlated SNPs via the 1000 Genomes EUR reference panel. A high threshold of an r^2^ value of 0.95 and within 100 kb was used to increase the number of SNPs available per gene, which in turn improved the ability to detect an effect. When linkage disequilibrium was present, we selected the SNP with the lowest *P* value. When clumping at r^2^ = 0.95, the SNPs are not strictly independent. This potential issue is remediated using experimental validation in a model organism. We used the inverse variance weighted method to model the effect of *KCNAB2* expression in neurons on AD risk and performed additional analyses using other model types. These sensitivity analyses include validating our results via a weighted median model, which is more robust for instrument variables. We also assessed the heterogeneity of the instrument variables via Cochran’s Q statistic, as well as their quality via the F statistic. Finally, we performed Egger regressions to assess pleiotropy.

### Statistical analyses

Statistical analyses were performed via R version 4.0.0 and GraphPad Prism (www.graphpad.com). The data are presented as the mean values, and the error bars indicate ± the SDs. The number of biological replicates per experimental variable (n) is indicated in either the respective figure or the figure legend. The data distribution was verified via the Shapiro–Wilk test, and subsequent statistical testing was subsequently performed. No sample was excluded from the analysis unless otherwise stated. Blinding was not performed. Significance is indicated as * for *P* ≦ 0.05, ** for *P* ≦ 0.01, *** for *P* ≦ 0.001, **** for *P* ≦ 0.0001 and NS for *P* ≥ 0.05.

### Digital image processing

The fluorescence images were acquired as uncompressed bitmapped digital data (TIFF format) and processed via Adobe Photoshop with established scientific imaging workflows. To visualise the pixel intensity, confocal images acquired with identical settings were processed via a five-tone heatmap in ImageJ.

## Code availability

All the code is available in our GitHub repository (https://yizhouyu.com/abeta_sleep).

## Data availability

All data is available in GitHub (https://yizhouyu.com/abeta_sleep). Access to the UK Biobank data can be applied for via the UK Biobank website (www.ukbiobank.ac.uk).

## Abbreviations

AD: Alzheimer’s disease
Hk: hyperkinetic
OXPHOS: oxidative phosphorylation

## Supplementary information

**Supplementary Table 1. Heatmap of statistically significant biochemical alterations in flies expressing Aβ42 or Aβ42Arc.**

The shaded cells indicate p ≤ 0.05 (red indicates that the mean values are significantly greater for that comparison; green values are significantly lower). The bolded blue text indicates 0.05 ≤ p ≤ 0.10. All the data were normalised to the Bradford protein assay measurements. This table is related to Figure 3. This table is also available here: https://github.com/izu0421/abeta_sleep/blob/main/Supplementary_Table_1.xlsx.

**Supplementary Table 2. Proteomic changes in Aβ42 or Aβ42Arc-expressing flies and controls.**

The fold changes are calculated based on individual quantitation levels of normalised log-transformed (base 2) abundance values from the mass spectrometer. The statistical tests used were linear models with variances moderated by the empirical Bayes method and peptide-spectrum match. The Adj.P.Val corresponds to a P value corrected via the Benjamini– Hochberg method. This table is related to Figure 5. This table is also available here: https://github.com/izu0421/abeta_sleep/blob/main/Supplementary_Table_2.xlsx.

